# Mechano-Biochemical Regulation of the C. elegans HMP1–HMP2 protein complex

**DOI:** 10.1101/2021.09.27.462086

**Authors:** Shimin Le, Miao Yu, Sterling Martin, Jeff Hardin, Jie Yan

## Abstract

The HMP1-HMP2 protein complex, a counterpart of *α*-catenin–*β*-catenin complex in C. elegans, mediates the tension transmission between HMR1 (cadherin) and actin cytoskeleton and serves as a critical mechanosensor at the cell–cell adherens junction. The complex has been shown to play critical roles in embryonic development and tissue integrity in C. elegans. The complex is subject to tension due to internal actomyosin contractility and external mechanical micro-environmental perturbations. However, how tension regulates the stability and interaction of HMP1–HMP2 complex has yet to be investigated. Here, we directly quantify the mechanical stability of the full-length HMP1 and its force-bearing modulation domains (M1-M3), and show that they unfold within physiological level of tension (pico-newton scale). The inter-domain interactions within the modulation domain leads to strong mechanical stabilization of M1 in HMP1, resulting in a significantly stronger force threshold to expose the buried vinculin binding site compared to the M1 domain in *α*-catenins. Moreover, we also quantify the mechanical stability of the inter-molecular HMP1–HMP2 interface and show that it is mechanically stable enough to support the tension-transmission and tension-sensing of the HMP1 modulation domains. Further, we show that single-residue phosphomimetic mutation (Y69E) on HMP2 weakens the mechanical stability of the HMP1–HMP2 interface and thus weakens the force-transmission molecular linkage and the associated mechanosensing functions. Together, these results provide a mechano-biochemical understanding of C. elegans HMP1–HMP2 protein complex’s roles in mechanotransduction.

## I. INTRODUCTION

Cell-cell adhesions play crucial roles in embryonic development and tissue integrity. These adhesions are formed and regulated by a variety of force-bearing proteins that are assembled into supramolecular physical linkages. The force-bearing proteins within the linkages interact with numerous signalling proteins in a manner sensitive to mechanical cues, forming the physical basis for mechanotransduction [1]. For instance, the classic cadherin-based adherens junction is linked to actin cytoskeleton through a core supramolecular physical linkage comprising of a linear array of cadherin, *β*-catenin, *α*-catenin and F-actin. The tension in the linkage has been estimated to be in the order of several piconewtons (pN) by molecular tension sensors [2–4]. Signalling proteins interact with mechanosensing proteins in the linkage in a force-dependent manner (e.g., vinculin–*α*-catenin interaction), regulating the strength of the adherens junction and converting the mechanical cues into a cascade of downstream biochemical signalling events [5–12]. While the mechano-biochemical regulations of the mammalian adherens junction have been intensively investigated over recent years [4, 6–12], its counterpart in C. elegans, an important model system for development and aging, has been much less understood [2–4].

In C. elegans, the adherens junction is linked to actin cytoskeleton through a core HMR1(cadherin)–HMP2(*β*-catenin)–HMP1(*α*-catenin)–F-actin supramolecular linkage [13–20]. The extra-cellular domains of HMR1 of one cell dimerizes with the HMR1 of the neighboring cells. This supramolecular linkage is subject to tensile forces due to internal actomyosin contraction and external mechanical microenvironmental perturbations, and is believed to play crucial mechanotransduction functions of cells in tissues [14, 17, 21]. However, how these core proteins respond to physiological level of tensions, are still unknown. Furthermore, the mechanical stability of the protein-protein interfaces in the linkage, which determines the integrity of the tension-transmission supramolecular linkage, has yet to be investigated.

In this study, we focused on the mechano-biochemical regulations of the HMP1 protein, and the protein-protein interface it forms with HMP2. By single-protein manipulation using magnetic-tweezer [22–24], we directly quantified the mechanical stability of the full-length HMP1 and its force-bearing modulation domains (M1-M3), as well as the HMP1–HMP2 interface. We showed that the HMP1 force-bearing domains unfold within pico-newton (pN) scale tensions. Interestingly and importantly, the inter-domain interaction between M1 and M2-M3 leads to strong mechanical stabilization of the M1 domain in HMP1, preventing high affinity binding of vinculin to the cryptic vinculin binding site (VBS) in the M1 domain during short exposruee to low tensions (< 15 pN), in sharp contrast to mammalian *α*-catenins where the VBSs are readily exposed at tension slightly above 5 pN [6, 12]. Furthermore, our results show that the HMP1–HMP2 interface has an average lifetime of tens of seconds to tens of minutes within physiologically relevant tension range. Finally, phosphotyrosine-mimetic mutations (Y69E) on HMP2 was shown to significantly weaken the mechanical stability of the HMP1–HMP2 interface. Together, these results reveal an intricate interplay between the mechanical and biochemical regulations of the HMR1–HMP2–HMP1 mediated force transmission and tension-dependent interactions, providing deeper insights into the molecular mechanisms underlying mechanotransduction at the adherens junction of C. elegans.

## II. RESULTS

### A. Mechanical stability of full-length HMP1

In order to probe the mechanical stability of full-length HMP1, we prepared a singlemolecule construct where the full-length HMP1 is spanned between four well-characterized titin Ig 27th domain (I27, two repeats at each end) acting as a molecular spacer [25–27]. The N- and C-termini of the construct contain a biotinylated AviTag and a SpyTag [26, 28], respectively, for specific tethering (Figure 1a). Furthermore, there is a 572-bp DNA handle added between the single-molecule construct and the super-paramagnetic bead as an additional spacer to avoid non-specific interaction between the bead and surface [26, 27, 29]. A magnetic-tweezer setup was implemented to apply well-controlled force on the tethered molecule via the bead and record the bead height in real time [22–24]. Due to force balance, the tension in the molecule is the same as the applied force. More details of the singlemolecule construct can be found in Texts S1.

**Figure 1.**
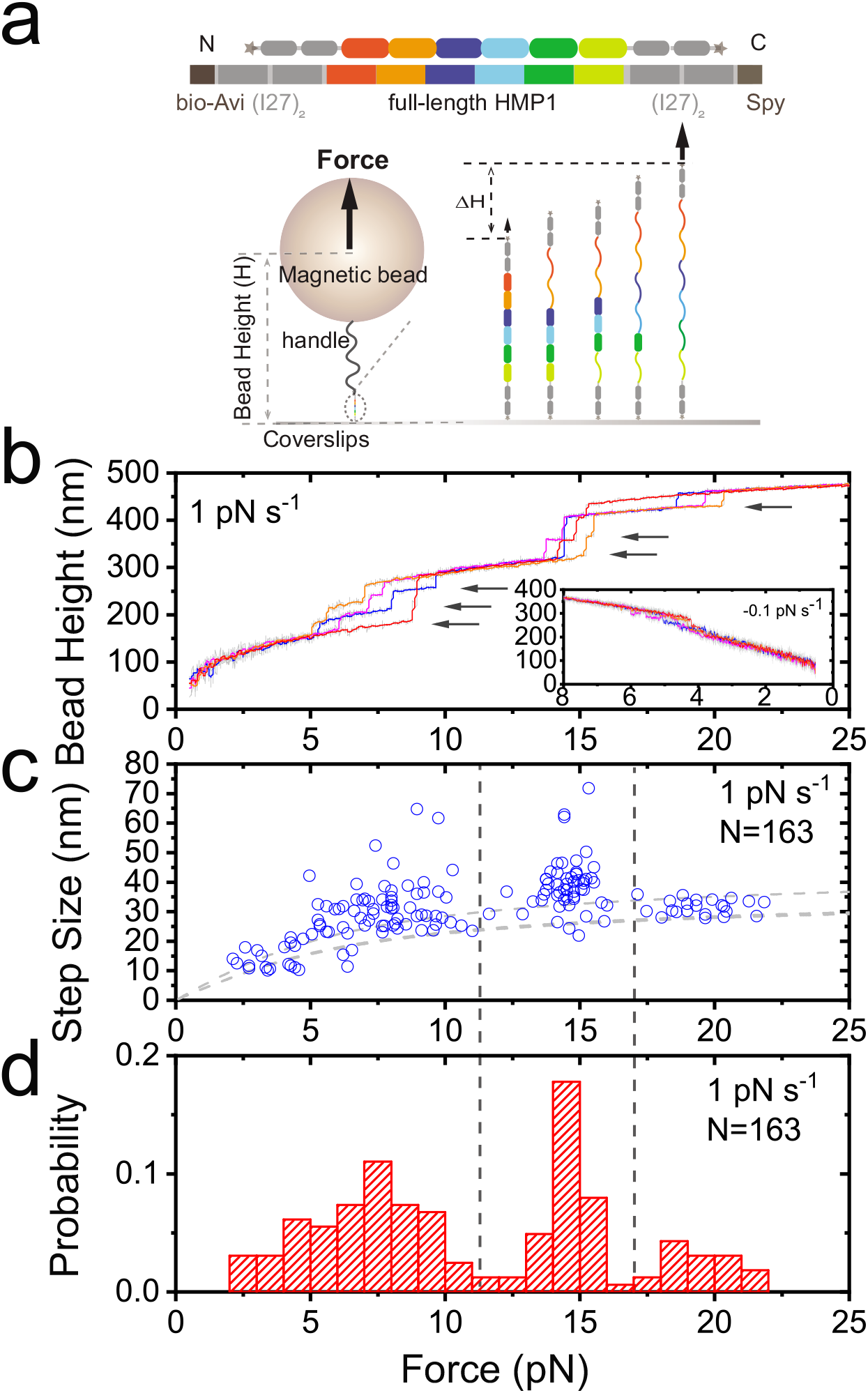
Mechanical stability of the full-length HMP1. (**a**). Sketches of the single-molecule construct and the experimental designs. Top panel shows the domain map of the construct: AviTag, two repeats of I27 domain (I27)_2_, full-length HMP1, (I27)_2_, SpyTag. The bottom panel shows a single-molecule tether between a SpyCather-coated coverslip surface and a biotin-DNA-coated super-paramagnetic bead (via streptavidin). Structural changes of the domains in the tether lead to extension changes of the molecule, which can be detected by the corresponding bead height changes. (**b**). Four representative force–height curves of a full-length HMP1 tether during force-increase scans at a loading rate of 1 pN s^-1^. The coloured curves are obtained by 10-point FFT (fast Fourier transformation, FFT) smooth of the raw data (grey). The arrows indicate the unfolding events of the six domains in HMP1. The inset shows the refolding events during force-decrease scans at a loading rate of −0.1 pN s^-1^. (**c-d**). The resulting force-dependent unfolding step sizes and the normalized force histogram obtained over 30 repeats of scans from three independent tethers. *N* in panel (d) indicates the total number of unfolding events.

Figure 1b shows the typical force–height curves of the full-length HMP1 during linear force-increase scans with a force-loading rate of 1 pN s^-1^ from ~ 1 pN up to 30 pN. During each force-increase scan, six unfolding events were clearly observed, indicated by six stepwise bead height jumps (black arrows in Figure 1b). Such stepwise bead height change equals the extension change of the tethered molecule due to structural changes [22, 30]. The I27 domain is highly mechanically stable, which unfolds with a slow rate of approximately 10^-3^ s^-1^ over the force range at lab room temperatures [25]. Therefore, these six unfolding events correspond to the unfolding of the six structured domains in full-length HMP1, i.e., two N-terminal domains (N1, and N2), three modulation domains (M1, M2 and M3), and the C-terminal F-actin binding domain. These unfolded domains could refold at forces <6 pN during force-decrease scans (inset of Figure 1b, with a loading rate of −0.1 pN ^-1^), allowing repeating the force scans for multiple cycles on one tethered molecule.

Performing over 30 force-increase scans, we obtained the force-step size graph of the unfolding events (Figure 1c) and the corresponding unfolding force distributions (Figure 1d). Clearly, mechanical unfolding of the six domains can be roughly categorized into three mechanical groups: three domains unfold at a force range of 3-10 pN, two domains unfold at 10-15 pN, and one domain unfolds at 16-22 pN. In addition, occasionally, two domains unfolded concurrently (within our temporal resolution: 10 ms), indicated by a single unfolding event associated with a step size that corresponds to the sum of the step size of two individual domains. The results from the mechanical unfolding of the full-length HMP1 suggest that all the functional domains of HMP1 respond to physiological level of forces [2–4].

### B. Mechanical stability of modulation domain of HMP1

As the modulation domains contain key binding sites for signalling proteins such as vinculin and SGRP1[18], we sought to probe the mechanical stability of the tension-bearing modulation domain of HMP1. Similar to the full-length HMP1 construct, we prepared a single-molecule construct “HMP1-M123” that contains the three modulation domains (Figure 2a). Figure 2b shows the typical force–height curves of the HMP1-M123 during force-increase scans with a force loading rate of 1 pN s^-1^, where three unfolding events of the three modulation domains were observed. Repeating such force-increase scans for over 100 times, we obtained the 2D graph of force-dependent sizes (Figure 2c) and the unfolding force distributions of the three modulation domains (Figure 2d). Two of the domains unfolded almost concurrently at about 10-15 pN, while the other unfolded at a higher force of 15-20 pN. Comparing to the unfolding signals obtained from the full-length HMP1 (Figure 1), the unfolding of modulation domains fall into the second and third mechanical groups of the full-length HMP1. Hence, the two N-terminal domains (N1, and N2) and C-terminal domain of HMP1 should belong to the first mechanical group that unfold within 3-10 pN (with a loading rate of 1 pN s^-1^). Together with results of next sections, we can assign the first two unfolding events to M1 and another domain from M2 or M3, and the third unfolding event to M2 or M3. The three unfolded modulation domains could refold in a stepwise manner during force-decrease scans at forces of ~ 4 pN, 3 pN and below 2 pN, respectively, at a force loading rate of −0.1 pN s^-1^ (Figure 2e).

**Figure 2.**
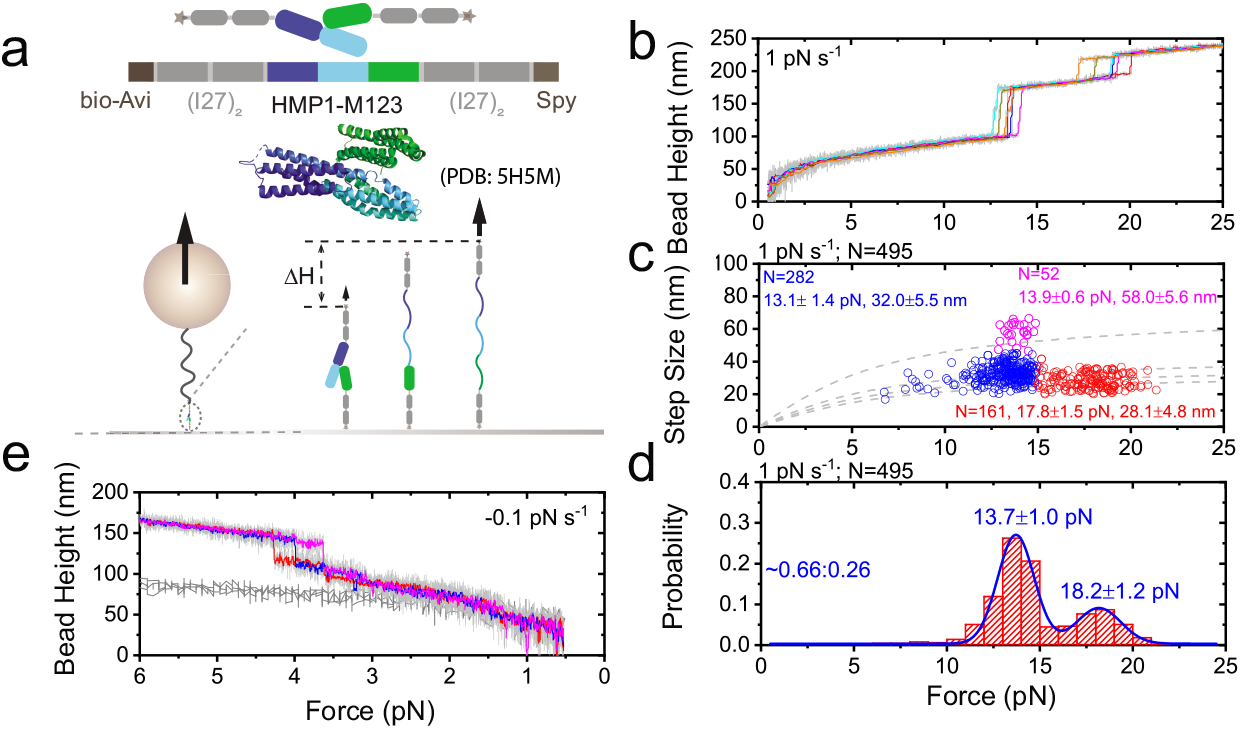
Mechanical stability of the HMP1 modulation domains. (**a**). Sketches of the single-molecule construct and the experimental design. Top panel shows the domain map of the construct, and the structure of the modulation domains (PDB:) [18]. The bottom panel shows a single-molecule tether under force. (**b**). Five representative force–height curves of the HMP1 modulation domains during force-increase scans at a loading rate of 1 pN s^-1^. (**c-d**). The resulting force-dependent step sizes and the normalized force histogram of the unfolding events obtained over 165 repeats of scans from five tethers. The unfolding events are divided into three groups based on unfolding forces and step sizes. The number of unflding events *(N*) in each group and the corresponding average unfolding forces and step sizes are indicated (panel (**c**)). The blue curve in panel (**d**) is the double-Gaussion fitting to the normalized unfolding force histogram. The peak forces are indicated. The area ratio (0.66:0.26) of the two groups is also indicated. (**e**). Three representative force–height curves (colored curves) of HMP1 modulation domains during forcedecrease scans with a loading rate of −0.1 pN s^-1^ started from fully unfolded conformation. As a comparison, the force–height curves of the HMP1 modulation domains with all three domain folded (gray curves) are also plotted.

Unexpectedly, the M1 domain of HMP1 exhibits a much higher mechanical stability than its mammalian counterparts such as *α*-catenins, where rapid unfolding of the M1 domain occurs at forces slightly above *~* 5 pN at similar force-loading rates [6, 12]. This significantly higher mechanical stability of the HMP1-M1 domain indicates either a stronger intrinsic mechanical stability of the M1 domain, or mechanical stabilization due to an interaction with its neighbouring domains.

### C. Tension-induced unfolding of HMP1 modulation domain activates high-affinity vinculin binding

Similar to mammalian *α*-catenin, there is a single conserved vinculin binding site (VBS) buried inside the HMP1-M1 domain, which has been shown to bind to DEB1 (vinculin counterpart in C. elegans), as well as mammalian vinculin[18]. Previous ITC (Isothermal titration calorimetry) experiments reported that the D1 domain of vinculin and DEB1 bind to HMP1-M123 domain with a low affinity – the vinculin-D1 binding was associated with a high dissociation constant of *K*_d_ ~ 600 nM, whereas the DEB1-D1 binding was not detectable [18]. On the other hand, both D1 domains bind to HMP1-M1 domain alone with a much higher affinity indicated by *K*_d_ of ~ 3 nM for vinculin-D1 and ~ 150 nM for DEB1-D1 [18]. These results suggest that the inter-domain interaction of M1–M23 greatly suppresses the binding of the D1 domain of vinculin or DEB1 to the VBS in the HMP1-M1 domain. Since HMP1 modulation domain lies in a tension transmission supramolecular linkage, it is reasonable to postulate that tension-induced disruption of M1–M23 interaction could release the auto-inhibitory inter-domain interaction, which in turn activates a high-affinity binding of vinculin/DEB1 to HMP1-M1.

To test this hypothesis, we first stretched the single-molecule construct of HMP1-M123 with multiple linear force-increase scans (1 pN s^-1^) followed by force-decrease scans (−0.1 pN s^-1^), and observed the characteristic cooperative unfolding of two modulation domains, one of which is M1, at 10-15 pN and the unfolding of the other modulation domain (M2 or M3) at 15-20 pN (Figure 3a, dark gray curves), as well as the characteristic refolding of the domains at lower forces < 5 pN (Figure 3e). We then introduced 10 nM vinculin-D1 (the vinculin-D1 domain, the binding domain for the VBS in M1) at low forces of ~ 1 pN (to ensure that the domains were folded during solution exchange), and repeated the force-increase and force-decrease scans (Figure 3a, magenta and orange curves). During the initial several force-increase scans, the characteristic unfolding signals of the three M domains were still observed (magenta curves), followed by losing the unfolding signature of one of domains at 10-15 pN (orange curve in Figure 3a and Figure 3f), which only occurred in the presence of vinculin-D1.

**Figure 3.**
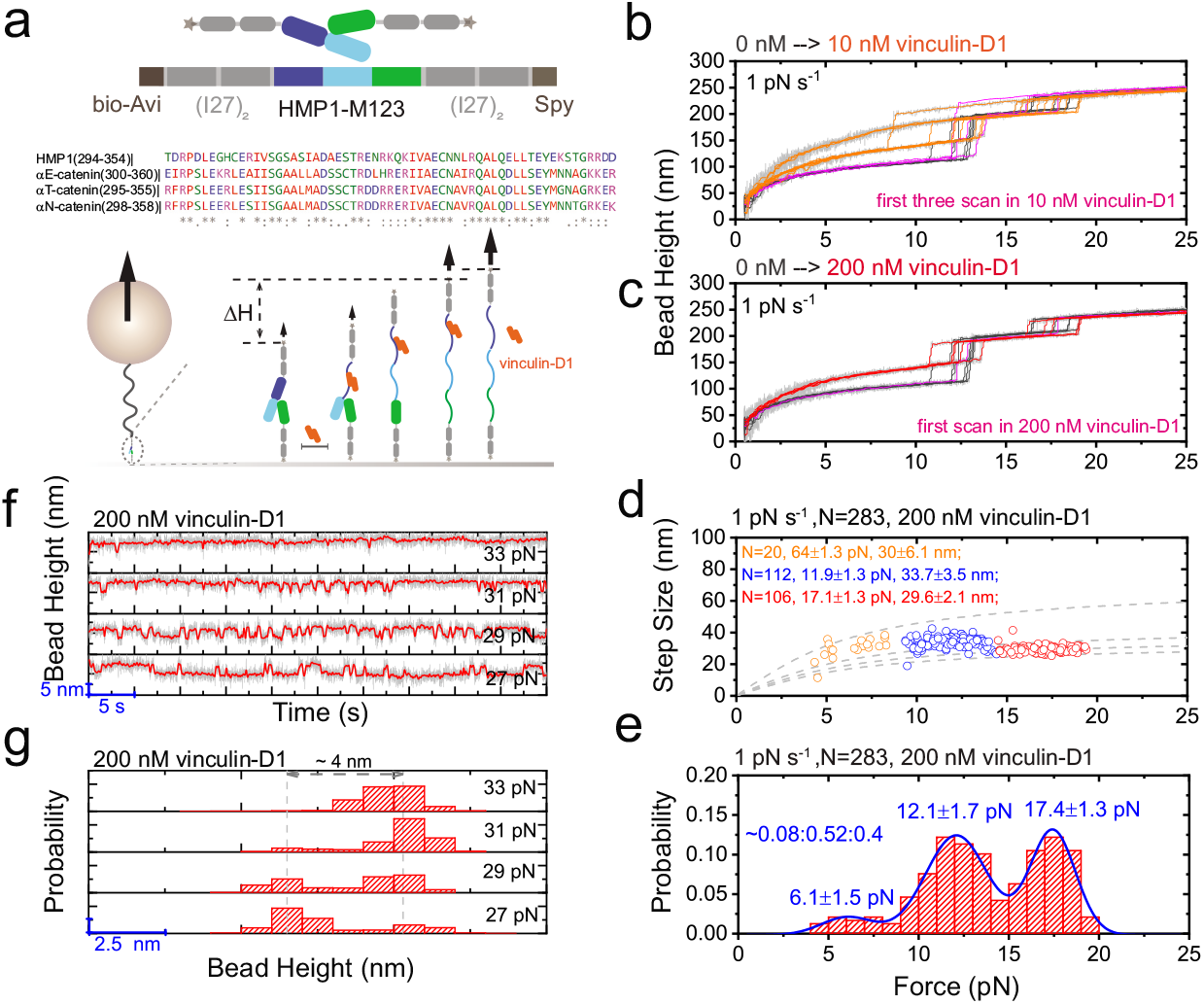
Mechanical activation of HMP1 modulation domains for vinculin binding. (**a**). The Left panel shows the domain map of the HMP1 modulation domain, and the sequence alignment of the conserved vinculin binding sites among the HMP1 and *α*-catenins. The right panel shows a single-molecule tether under force. (**b**). Representative force–height curves during force–increase scans in the absence of vinculin-D1 (dark grey curves) and in the presence of 10 nM vinculin-D1 (magenta and orange curves). (**c**). Representative force–height curves during force-increase scans in the absence of vinculin-D1 (dark grey curves) and in the presence of 200 nM vinculin-D1 (magenta and red curves). (**d-e**). The resulting force-dependent step sizes and the normalized unfolding force histogram obtained over 150 repeats of scans from five tethers in 200 nM vinculin-D1. The unfolding events are divided into three groups based on unfolding forces. The number of unfolding events (N) in each group and the corresponding average unfolding forces and step sizes are indicated (panel (**d**)). The blue curve in panel (**e**) is the triple-Gaussion fitting to the normalized unfolding force histogram. The peak forces and the area ratio (0.08:0.52:0.40) of the three groups areindicated. (**f**) Representative force-height curves of the tether at indicated constant forces. Dynamic binding or unbinding events of vinculin-D1 were indicated by 3-5 nm bead height changes, as shown in the resulting normalized histograms of bead height (**g**).

The observation of vinculin-D1 binding at 10 nM under force is in sharp contrast to the *K*_d_ ≈ 600 nM measured between vinculin-D1 and HMP1-M12 in the absence of force, consistent with the hypothesis that the force-induced unfolding of HMP1 modulation domains exposes the buried VBS for high-affinity vinculin-D1 binding. The high-affinity vinculin-D1 binding then suppresses the refolding of the M1 domain during the force-decrease scans. At a higher concentration (200 nM) vinculin-D1 (Figure 3c), the binding of vinculin-D1 was observed after the unfolding of the M domains in the first force-increase scan, indicating faster binding at the higher concentration. Figure 3d-e show the resulting force-dependent step sizes and transition forces of the remaining M2 and M3 domain unfolding events in the presence of 200 nM vinculin-D1 in solution. The unfolding force histogram show two major mechanical groups spread over 10-15 pN and 15-50 pN force ranges, respectively. Compared to the unfolding force distribution in the absence of vinculin-D1 (Figure2c-d), the HMP1-M1 refolding is suppressed by vinculin-D1 binding, while the refolding and unfolding signatures of HMP1-M2 and HMP1-M3 were largely unaffected. There is a minor mechanical group with unfolding forces less than 10 pN, which might be due to an altered refolding of the modulation domains due to vinculin-D1 binding.

In addition, the vinculin-D1 binding to VBS is known to induce a *α*-helical structure of the VBS peptide associated with a different force-dependent extension from the randomly coiled conformation [6, 12, 26], which implies that the applied force can tune the binding/unbinding dynamics of the vinculin-D1 to VBS. Indeed, when the force was increased to a range of 27 — 30 pN in the presence of 200 nM vinculin-D1, dynamic transitions between two distinct extension levels were observed, associated with an extension difference of ~ 4 nm (Figure 3g-h), which is the expected extension difference between the *α*-helical and randomly coiled conformations of the VBS peptide over this force range [6, 12, 26]. This result suggests that the bound complex of VBS–vinculin-D1 can be rapidly destabilized at forces greater than this force range.

### D. HMP1–HMP2 interface provides sufficient mechanical stability for supporting tension transmission

We have shown that the tension-induced rapid unfolding of modulation domains involves tensions in the range of 10-15 pN. Tension-dependent activation of trans-binding of signalling proteins requires that the tension-transmission HMR1–HMP2–HMP1–F-actin supramolecular linkage can withstand this tension range over a sufficient lifetime to support the tension dependent interactions. As the integrity of the tension-transmission pathway is dependent on the mechanical stability of protein–protein interfaces in the linkage, it is important to measure the mechanical stability of protein–protein interfaces in the tension transmission linkage. The HMP1–HMP2 interface is a critical protein–protein interface in the linkage, which comprises of the N-terminal tail peptide of HMP2 (Nt) and the N-terminal domain of HMP1 (N12) (Figure 4a) [19]. We sought to quantify the mechanical stability of this interface.

**Figure 4.**
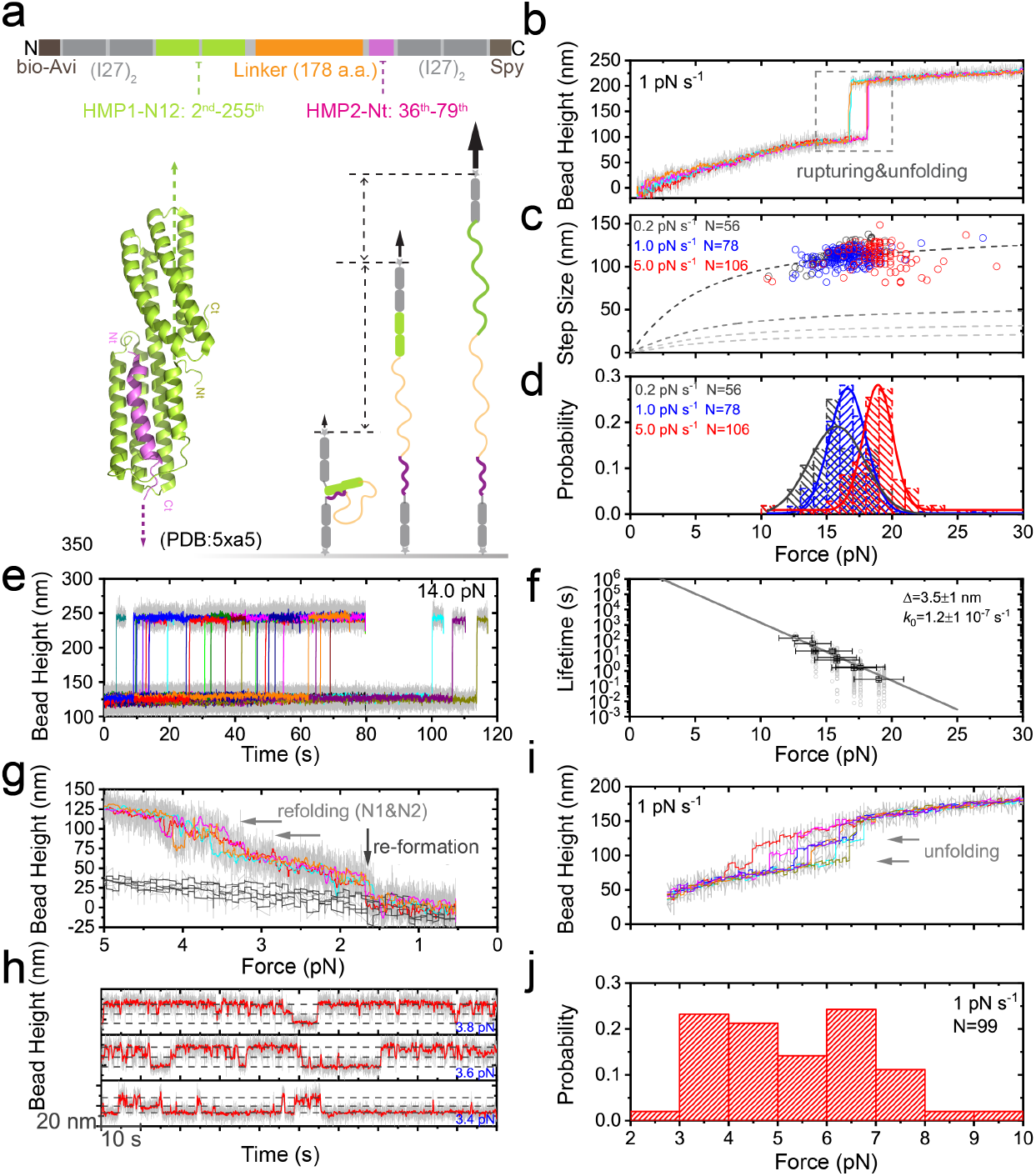
Mechanical stability of the HMP1–HMP2 interface. (**a**). The single-molecule construct, the domain map, the structure of the HMP1–HMP2 interface (PDB:) [19], and the single-molecule tether under force. (**b**). Representative force–height curves of a tether during force-increase scans at a loading rate of 1 pN s^-1^. (**c-d**). The resulting force-dependent step sizes and the normalized rupturing force histograms obtained over 50 repeats of scans for five tethers at three indicated force loading rates. The number of rupturing events, the average forces and the average step sizes at corresponding loading rates are *N*=56, *F* =15.9±1.9 pN, Δ*H* =115.4±12.2 nm (0.2 pN s^-1^); *N*=178, *F*=16.58±1.7 pN, Δ*H*=110.9±8.4 nm (1 pN s^-1^); *N*=106, *F*=19.1±3.1 pN, Δ*H* =112.6±14.2 nm (5 pN s^-1^). (**e**). Representative force–height curves of a tether during forceclamp at 14.0 pN. The stepwise bead height jumps indicate the rupturing of the HMP1–HMP2 interface. (**f**). The resulting force dependent average lifetime *τ(F*) of the HMP1–HMP2 interface obtained from such force-clamp experiments, which can be well fitted with Bell’s model, 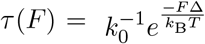, where Δ is the transition distance and *k*_0_ is the extrapolated zero-force rupturing rate. (**g**). Representative force–height curves of the tether during force-decrease scans with a loading rate of −0.1 pN s^-1^. Grey arrows indicate the refolding of N1 and N2 domains, black arrow indicates the re-formation of the interface. (**h**). Representative time traces of the bead height at indicated constant forces, before the re-formation of the interface. Dynamic unfolding and refolding of the N1 and N2 domain were observed. (**i**). Representative force–height curves of the tether during forceincrease scans at the indicated loading rate starting from a conformation where the HMP1–HMP2 interface was not re-formed, whereas the N1 and N2 domains were refolded. (**j**). The resulting normalized unfolding force histogram during the force-increase scans.

We utilized a single-molecule construct that converts a protein–protein interaction into an intra-molecular interaction (Figure 4a) [10, 11, 26, 32–34]. Briefly, the construct of the HMP1–HMP2 interface consists of the N12 domain of HMP1, the Nt peptide of HMP2, and a long flexible unstructured peptide linker in between. The construct is spanned between two repeats of I27 domains at each side. An AviTag and a SpyTag are fused at each end of the construct for specific tethering (Figure 4a, and Text S1). At sufficiently low forces, N12 and Nt can interact with each other to form the HMP1–HMP2 interface, looping the long linker inside as a result. In contrast, a sufficiently high force can destabilize the interface, and the resulting rupturing of the interface leads to the release of the long linker. Such rupturing event leads to an abrupt extension increase of the construct, which is indicated by a stepwise large bead height increase of the same amount as the extension increase [30].

Figure 4b shows the typical force–height curves of the HMP1–HMP2 interface construct during force-increase scans with a loading rate of 1 pN s^-1^. Clearly, during each scan, a single bead height jump of 110.9±8.4 nm was observed at 16.6± 1.7 pN. The size of the bead height increases at these forces are consistent with the extension of the linker and unfolding of the domain N12 domain, suggesting concurrent rupturing of the interface and the unfolding of the N12 domain. Similar force-increase scans were repeated for over 50 times at three force-loading rates: 0.2 pN s^-1^, 1 pN s^-1^, 5 pN s^-1^, and the resulting force-dependent step sizes of the rupturing events are shown in Figure4c. The normalized histograms of rupturing forces at the three loading rates show that the interface ruptures at 15.9±1.9 pN, 16.58±1.7 pN, and 19.1±3.1 pN, respectively. These forces are greater than the unfolding forces of the first and second modulation domains at the corresponding loading rates, suggesting that the HMP1–HMP2 interface provides a mechanical stability that is enough to support the mechanotransduction of the modulation domains that require mechanical destabilization of the M1 domain for high affinity vinculin binding.

To directly quantify the tension-dependent lifetime of the HMP1–HMP2 interface, we implemented a force-clamp assay. Briefly, the construct was firstly held at a force of ~ 0.5 pN for 30 sec to allow the formation of the interface, and then held at a targeting force of 12-16 pN and recorded the dwell time till the interface was ruptured. Repeating the procedure for multiple cycles (> 50), the average lifetime of the interface at this force was obtained by exponential fitting to the histogram of the dwell times (Methods). The force-dependent lifetimes of the interface were obtained by performing such measurement at different targeting forces (Figure 4e-f). The force-dependent lifetime of the interface consistently shows that the interface is more mechanically stable than the M1. For an example, at the force of ~ 14 pN the HMP1–HMP2 interface can last over tens of seconds, while at the same force the M1 domains unfold almost immediately.

In addition, the construct also allowed us to quantify the mechanical responses of N12 domains of HMP1. Consistent with the data from full-length HMP1 unfolding experiments, the N1 and N2 domain unfolded at forces of 3-9 pN during force-increase scans at 1 pN s^-1^ loading rate (Figure 4g). The equilibrium unfolding/refolding dynamics of N1 and N2 domains can be observed at constant forces around 4 pN (Figure 4h).

### E. Single-residue phosphorylation-mimetic mutations on HMP2 weaken the mechanical stability of the HMP1–HMP2 interface

The HMP2 N-terminus peptide contains two phosphorylation residues, i.e., the 69_th_ tyrosine (Y69) and 47_th_ serine (S47) residues, which lies within the HMP1–HMP2 interface. It was previously observed that the Y69E or S47E mutation of the HMP2 led to cell–cell adhesion disruptions [19]. This has led to a hypothesis that the phosphorylation of these residues could weaken the stability of the HMP1–HMP2 interface, which has yet to be tested [19]. In this section, we quantify the effects of these mutations on the mechanical stability of the HMP1–HMP2 interface. For this purpose, we prepared two single-molecule constructs of the HMP1–HMP2 interface that carry the 1) Y69E mutation, or 2) S47E mutation (Figure 5a).

**Figure 5.**
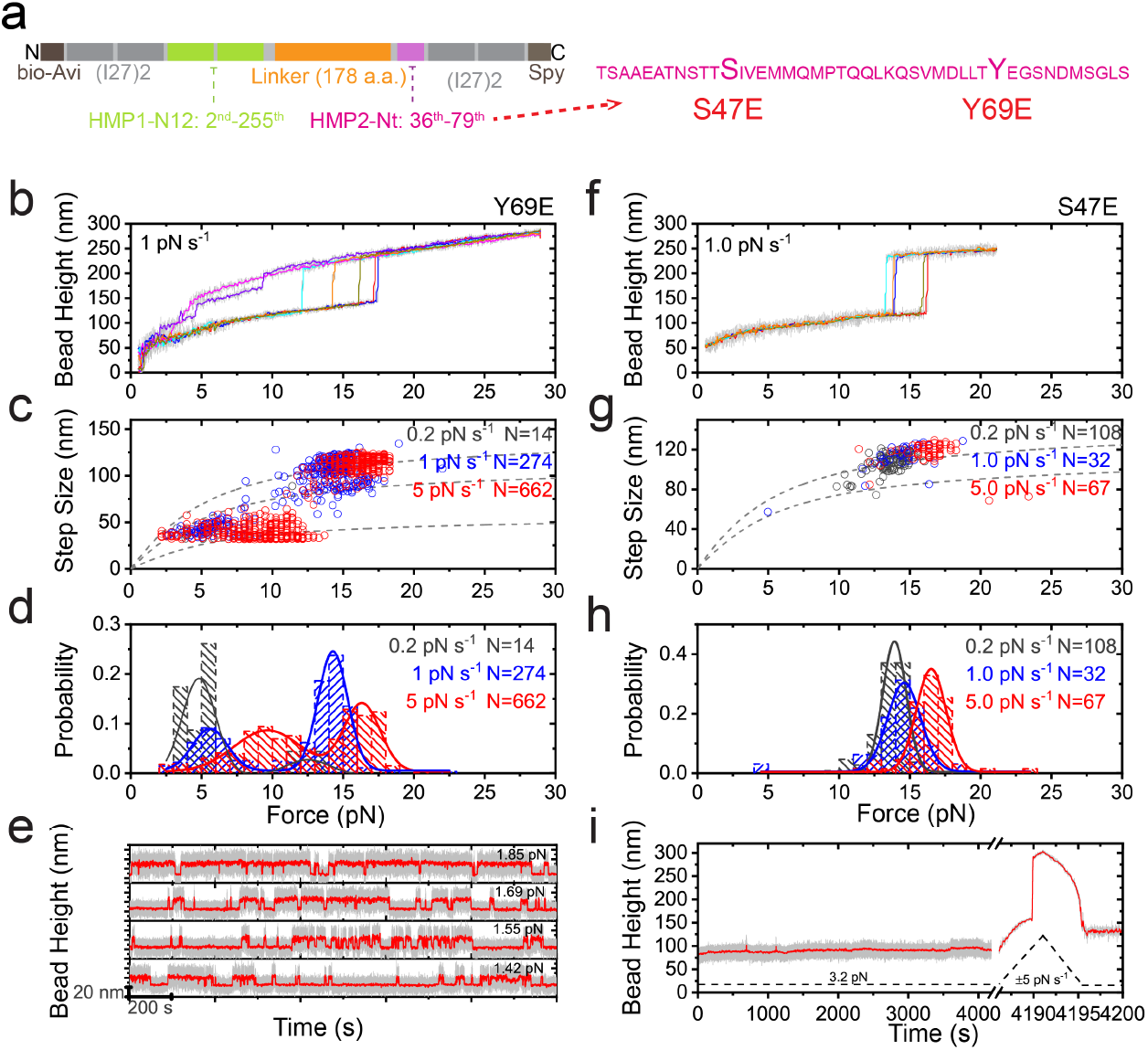
Mechanical stability of the HMP1–HMP2 interface with Y69E or S47E mutation on HMP2. (**a**). Sketches of the domain map and sequences of HMP2-Nt. (**b**). Representative force–height curves of a HMP1–HMP2^Y69E^ interface tether during force-increase scans at 1 pN s^-1^. (**c-d**). The resulting force-dependent step sizes and the normalized rupturing force histograms obtained at the three indicated loading rates. The number of rupturing events, the average forces and the average step sizes at corresponding loading rates are *N*=12, *F*=4.7±0.9 pN, Δ*H*=38.2±4.5 nm; *N*=2, *F*=12.6±0.7 pN, Δ*H* = 100.4±16.1 nm (0.2 pN s^-1^, two groups); *N*=86, *F*=5.7±1.5 pN, Δ*H*=44.5±9.3 nm; *N*=188, *F*=14.1±1.4 pN, Δ*H*=105.8±14.3 nm (1.0 pN s^-1^, two groups); *N*=221, *F*= 7.4±2.1 pN, Δ*H*=39.9±7.3 nm; *N*=441, *F*= 14.7±2.5 pN, Δ*H*=91.5±33.8 nm (5.0 pN s^-1^, two groups); (**e**). Representative time traces of the bead height of a HMP1–HMP2^Y69E^ interface tether at indicated constant forces, where dynamic formation and rupturing of the interface were observed. (**f**). Representative force–height curves of a HMP1–HMP2^S47E^ interface tether during force-increase scans at 1 pN s^-1^. (**g-h**). The resulting force-dependent step sizes and the normalized rupturing force histograms obtained at the three indicated loading rates. The number of rupturing events, the average forces and the average step sizes at corresponding loading rates are:*N*=108, *F*= 13.6±1.2 pN, Δ*H*=107.4±9.2 nm (0.2 pN s^-1^); *N*=32, *F*= 14.3±2.2 pN, Δ*H* =112.4±14.7 nm (1.0 pN s^-1^); *N*=67, *F*= 16.3±1.7 pN, Δ*H* =117.3±10.9 nm (5.0 pN s^-1^); (**i**). Representative time traces of the bead height of a HMP1–HMP2^S47E^ tether at a constant force of 3 pN for over 4000 sec where the interface remained stable, followed by a force-increase scan (after the break sign) to rupture the interface.

Figure 5b shows typical force–height curves of the HMP1–HMP2^Y69E^ interface construct during force-increase scans with a force loading rate of 1 pN s^-1^. Repeating similar force-increase scans for three force-loading rates: 0.2 pN s^-1^, 1 pN s^-1^, 5 pN s^-1^, we obtained the resulting force-dependent step sizes of the interface rupturing (corresponding to the first stepwise bead height increase) and domain unfolding transitions (Figure 5c), and the corresponding interface rupturing force distributions at three force-loading rates (Figure 5d). Interestingly, a two-peak distribution of the interface rupturing force was observed. For instance, at 1 pN s^-1^, one peaked at 14.1±1.4 pN, and the other peaked at 5.7±1.5 pN, which is in sharp contrast to the single-peaked rupturing force distribution (peak force at 16.58±1.7 pN for 1 pN s^-1^) observed from the wild-type interface. Similar experiments performed on the HMP1–HMP2^S47E^ interface shows a single-peak distribution (peak force at 14.3±2.2 pN), which is slightly lower than the peaked force of the wild-type interface (Figure 5e-g). These results demonstrate that the phosphorylation of the two residues lead to impaired mechanical stability of the HMP1–HMP2 interfaces. Between the two, the results suggest that the Y69E mutations could cause a more severe weakening effect on the mechanical stability of the interface. Consistently, the lifetimes of the HMP1–HMP2^Y69E^ interface were within 200 seconds at forces of 1-2 pN, while the HMP1–HMP2^S47E^ interface remained stable over thousands of seconds at a slightly higher force of ~ 3 pN.

## III. DISCUSSIONS

In this work, aiming to provide insights into the molecular mechanisms underlying the mechano-biochemicial regulation of the HMP1–HMP2 mediated cell–cell adhesions of C elegans, we have systematically investigated 1) the mechanical stability of the full-length HMP1 protein, particularly the functional modulation domains within physiological force range; 2) the force-dependent binding of vinculin to HMP1; 3) the mechanical stability of the HMP1–HMP2 interface and its dependence on the phosphorylation of the HMP2 resides within the interface.

Our results revealed that the five domains within HMP1 can be unfolded individually within physiological level of forces less than 20 pN. An interesting finding is that, despite the high-degree of structural and domain organization similarities between HMP1 and its mammalian counterpart *α*-catenins, we observed significant differences in the mechanical stability of the modulation domains. Importantly, the unfolding force of the critical VBS-bearing M1 domain is shifted from ~ 5 pN for mammalian *α*-catenins to ~ 14 pN for HMP1, revealing a significantly greater autoinhibition of M1 unfolding, and hence its binding to vinculin, by its inter-domain interaction between M1 and M2-M3. A previous structural study has shown inter-domain interactions of M1–M23 in HMP1 [18]; however, whether it can lead to the observed mechanical stabilization is unclear, since similar inter-domain interactions are also observed in mammalian *α*-catenins [31].

We show that the force-induced unfolding of HMP1-M1 is required for activation of high-affinity vinculin binding. Force-induced exposure of VBSs for high-affinity vinculin binding was observed for other VBS-containing force-bearing mechanosensing proteins, such as talin, *α*-actinin [6, 26]. Therefore, force-activated binding of vinculin seems a conserved strategy utilized by cells in different organisms.

Importantly, we show that the HMP1–HMP2 interface has a significantly greater mechanical stability than that of M1 and M2 domains, suggesting that this interface is mechanically stable enough to support the force-dependent interaction between the modulation domains and signalling factors such as vinculin. We also demonstrated that the mechanical stability of the HMP1–HMP2 interface is weakened by single phosphomimetic (Y69E, S47E) mutations of HMP2, implying destabilization of the tension-transmission supramolecular linkage and thus suppression of the activation of the modulation domains for their respective interactors. These results suggest the existence of an intricate interplay between mechanical and biochemical regulations of the HMP1–HMP2 mediated mechanosensing at cell-cell adhesions in C. elegans, which is consistent with the previous observation that phosphomimetic mutation on HMP2 led to adherens junction disruption in vivo [19].

An interesting question is raised regarding the molecular mechanisms that confer the HMP1–HMP2 interface the high mechanical stability. The structure of the HMP1–HMP2 interface reveals that the interface is formed by interactions between a single *α*-helix from the N-terminus of HMP2 and the N-terminal 4-*α*-helical bundle (N1 domain) of HMP1, leading to a 5-*α*-helical bundle in the complex. Actomyosin contraction generates tensile forces in the tension-transmission linkage, as a result the HMP1–HMP2 interface is under a shear-force pulling geometry (i.e., the force direction is nearly parallel to the interacting interface). Such protein-protein interface arrangement and force pulling geometry have also been found in several other critical force-bearing interfaces, such as the vinculin– talin, vinculin–*α*-catenin, *β*-catenin–*α*-catenin, which lie in different tension-transmission supramolecular linkages playing critical mechanotransduction roles at focal adhesion and cell-cell adhesions [1]. Together, these results suggest that protein-protein interfaces formed between a peptide and a rigid structural domain under a shear-force pulling geometry could be a evolutionarily conserved key mechanism to provide high mechanical stability to these tension-bearing interfaces [1].

## IV. METHODS AND MATERIALS

### Plasmids constructs and protein expression

Five plasmids were prepared for expression of the protein constructs for single-molecule stretching experiments: 1). pET151-Avi-(I27)_2_-(full length HMP1)-(I27)_2_-Spy, 2) pET151-Avi-(I27)_2_-(HMP1-M123)-(I27)_2_-Spy, 3) pET151-Avi-(I27)_2_-(HMP1-N12)-(long flexible linker)-(HMP2-Nt)-(I27)_2_-Spy, 4) pET151-Avi-(I27)_2_-(HMP1-N12)-(long flexible linker)-(HMP2-Nt^Y69E^)-(I27)_2_-Spy, 5) pET151-Avi-(I27)_2_-(HMP1-N12)-(long flexible linker)-(HMP2-Nt^S47E^)-(I27)_2_-Spy. Briefly, the DNA fragments encoding HMP1 domains were amplified by PCR the template sequence [18, 19]. The long flexible linker and other DNA fragments were synthesized by geneArt/IDTgblock. The DNA fragments were then assembled into a pET151-avi-(I27)_2_-(inset)-(I27)_2_-spy plasmid template [26] by HiFi DNA Assembly (NEB) and sequencing-confirmed (1st BASE). Each plasmid was co-transformed with a BirA plasmid and expressed in Escherichia coli BL21 (DE3) cultured in LB-media with D-Biotin (Sigma Aldrich), and affinity purified through 6His-tag. Detailed sequence information of the plasmids can be found in Text S1.

### Single-protein manipulation and analysis

A vertical magnetic tweezers setup was combined with a disturbance-free, rapid solution-exchange flow channel for conducting in vitro protein stretching experiments [22–24]. All in vitro protein stretching experiments were performed in solution containing: 1X PBS, 1% BSA, 2 mM DTT, 10 mM sodium L-ascorbate at 22 ± 1°C. The force calibration of the magnetic tweezers setup has a 10% uncertainty due to the heterogeneity of the diameter of paramagnetic beads [22] and the bead height determination of the magnetic tweezers setup has a ~ 2-5 nm uncertainty due to the thermal fluctuation of the tethered bead and the resolution of the camera [10].

## V. AUTHOR CONTRIBUTIONS

S.L., M.Y., J.Y. and J.H. conceived the study. S.L. and M.Y. designed the single molecule assay, performed the experiments and analyzed the data; S.L., M.Y. and S.M. prepared the plasmids; S.L., M.Y., J.Y. and J.H. interpreted the data; S.L., M.Y. and J.Y. wrote the paper.

## VI. ACKNOWLEDGEMENT

The authors thank the protein expression core and science communication core of the Mechanobiology Institute. The research is funded by the Singapore Ministry of Education Academic Research Fund Tier 2 (MOE2019-T2-1-099 to J.Y.), the Ministry of Education under the Research Centres of Excellence programme (to J.Y.), and the National Research Foundation, Prime Minister’s Office, Singapore, under its NRF Investigatorship Programme (NRF Investigatorship Award NRF-NRFI2016-03 to JY).

## Supplementary Information

### SUPPLEMENTARY TEXTS

#### Supplementary Text S1. Plasmids construct and protein expression

Five plasmids were prepared for expression of the protein constructs for single-molecule stretching experiments: 1). pET151-Avi-(I27)_2_-(full length HMP1)-(I27)_2_-Spy, 2) pET151-Avi-(I27)_2_-(HMP1-M123)-(I27)_2_-Spy, 3). pET151-Avi-(I27)_2_-(HMP1-N12)-(long flexible linker)-(HMP2-Nt)-(I27)_2_-Spy, 4). pET151-Avi-(I27)_2_-(HMP1-N12)-(long flexible linker)-(HMP2-Nt^Y69E^)-(I27)_2_-Spy, 5). pET151-Avi-(I27)_2_-(HMP1-N12)-(longflexiblelinker)-(HMP2-Nt^S47E^)-(I27)_2_-Spy. Briefly, the DNA fragments encoding HMP1 domains were amplified by PCR the template sequence [1, 2]. The long flexible linker and other DNA fragments were synthesized by geneArt/IDTgblock. The DNA fragments were then assembled into a pET151-avi-(I27)_2_-(inset)-(I27)_2_-spy plasmid template [3] by HiFi DNA Assembly (NEB) and sequencing-confirmed (1st BASE). Each plasmid was co-transformed with a BirA plasmid and expressed in Escherichia coli BL21 (DE3) cultured in LB-media with D-Biotin (Sigma Aldrich), and affinity purified through 6His-tag (at N-terminus). Detailed sequence information of the plasmids are listed below:

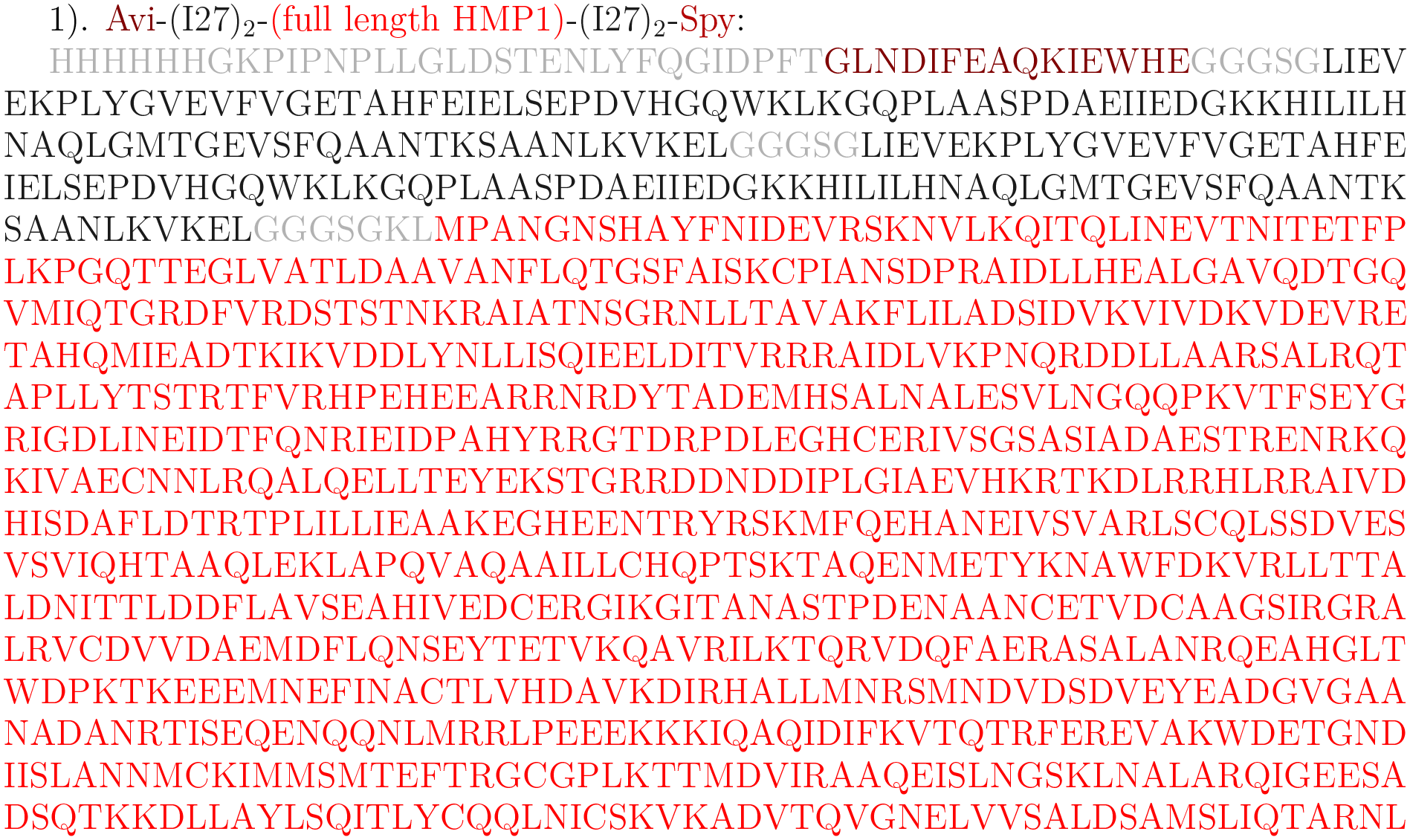
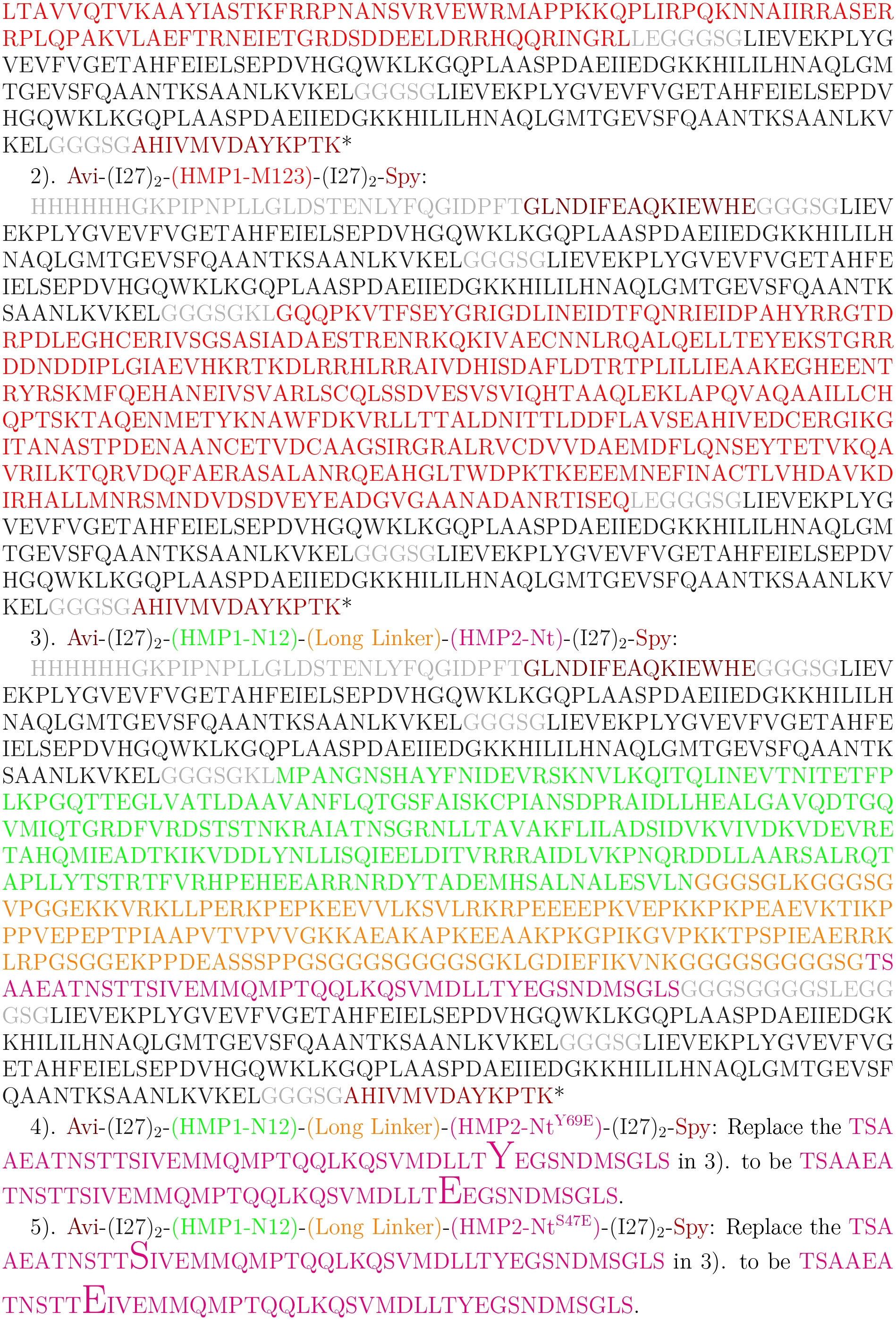

## Notes

### Competing Interest Statement

The authors have declared no competing interest.

